# The core effector RipE1 of *Ralstonia solanacearum* interacts with and cleaves Exo70B1 and is recognized by the Ptr1 immune receptor

**DOI:** 10.1101/2022.08.31.506019

**Authors:** Dimitra Tsakiri, Konstantinos Kotsaridis, Sotiris Marinos, Vassiliki A. Michalopoulou, Michael Kokkinidis, Panagiotis F. Sarris

## Abstract

*Ralstonia solanacearum* depends on numerous virulence factors, also known as effectors, to promote disease in a wide range of economically important host plants. Although some of these effectors have been characterized, none have yet been shown to target the host’s secretion machinery. Here, we used an extended library of NLR plant immune receptor integrated domains (IDs), to identify new effector targets. The screen uncovered that the core effector RipE1, of the *R. solanacearum* species complex, among other targets, associates with Arabidopsis exocyst component Exo70B1. RipE1, in accordance with its predicted cysteine protease activity, cleaves Exo70B1 *in vitro* and also promotes Exo70B1 degradation *in planta*. RipE1 enzymatic activity additionally results in the activation of TN2-dependent ectopic cell death. TN2 is an atypical NLR that has been proposed to guard Exo70B1. Despite the fact that RipE1 has been previously reported to activate defense responses in model plant species, we present here a *Nicotiana* species, in which RipE1 expression does not activate cell death. In addition, we discovered that RipE1 is recognized by Ptr1, a *Nicotiana benthamiana* CC-NLR, via its cysteine protease activity. Overall, this study uncovers a new RipE1 host target and a new RipE1-activated NLR while providing evidence and novel tools to advance in-depth studies of RipE1 and homologous effectors.

**Author Summary:** Bacterial wilt disease caused by *Ralstonia solanacearum*, poses a serious global threat for a wide range of agriculturally important plant species. This Gram-negative bacterium utilizes a collection of Type III Secretion System (T3SS) effectors to manipulate host cell defense and physiology. In this study, we searched for new subcellular plant targets of the core *R. solanacearum* effector RipE1, a cysteine protease. We discovered that RipE1 has multiple potential eukaryotic targets and further elucidated its association with the host exocyst complex. Using Artificial Intelligence (AI)-based predictions and performing both *in vitro* and *in planta* assays, we found that RipE1 promotes the degradation of plant exocyst component Exo70B1 through its enzymatic activity. Apart from being the first report of a *R. solanacearum* effector targeting a component of the host secretion machinery, our findings also identify an NLR from a model plant species that is able to recognize RipE1 protease activity and provide evidence that can lead to the discovery of additional RipE1 targets inside the host cell.

## Introduction

Plants, unlike animals, solely rely on their innate immune system to activate defense responses and fight off pathogens (1,2). To counter these responses, pathogens have evolved multiple virulence components, also known as effectors, that manipulate eukaryotic cell host metabolism, including pathways involved in innate immunity (3,4).

Plant innate immunity involves two intertwined cross-talking pathways that discern signs of invasion (5,6). Plant pattern-recognition receptors (PRRs) sense extracellular danger signals, mostly pathogen-associated molecular patterns (PAMPs), and activate a series of defense responses that constitute PAMP-triggered immunity (PTI) (7,8). Pathogen effectors translocated inside the host cell to block PTI, are recognized by plant intracellular nucleotide-binding leucine-rich repeat receptors (NLRs) that subsequently trigger the effector-triggered immunity (ETI) responses (7,9). Effectors’ recognition often leads to induction of a focused rapid programmed cell death known as the hypersensitive response (HR) that stops pathogen spread (10,11). NLRs can recognize effectors either by directly binding or indirectly by perceiving modifications caused by effectors’ activity on the operational target proteins (guardee model) or by a second protein that mimics their cellular target (decoy model) (12,13). A specialized version of direct recognition by NLRs is described via NLR integrated protein domains (NLR-IDs) (12,14). This model refers to NLRs carrying non-canonical domain fusions (integrated domains – IDs) that resemble pathogen targets to attract effectors and facilitate recognition (14,15). The evolutionary arms race between plants and microbial pathogens continues with either effector modification or the rise of new effectors that interfere with NLR activation or other downstream ETI signaling events, resulting in effectors’ triggered susceptibility (ETS) (16).

The exocyst is an octameric protein complex that mediates tethering of secretory vesicles to the plasma membrane, before their SNARE (soluble N-ethylmaleimide-sensitive factor attachment protein receptors) - dependent fusion (17,18). Although it was first identified in yeast, the exocyst complex is highly conserved among eukaryotes and is also present in plants (18–20). Among exocysts from different organisms that have been studied to date, the complex seems to be composed of the same eight subunits -Sec3, Sec5, Sec6, Sec8, Sec10, Sec15, Exo70, and Exo84-with the main difference in plants being the multiple paralogues encoding some of the subunits (20,21). In plants, the exocyst complex is involved in many physiological processes and there is strong evidence of its contribution to both PTI and ETI responses (22–29). The exocyst’s role in plant immunity is further validated through the fact that different pathogens have acquired effectors that target exocyst components or their interactors and compromise defense-related exocytosis (30– 35). In plants, Exo70 paralogues seem to be the most abundant, with Arabidopsis having 23 different Exo70 genes (24). Several of these Exo70 paralogues have been implemented in plant immune responses and are required for defense against distinct pathogens (25, 36, 37). Specifically, Arabidopsis Exo70B1 role in plant defense is highlighted by not only its interaction with key regulators and immune receptors, but also, by its involvement in plant autophagy, an emerging player in plant immunity responses (38–42). Furthermore, a number of different pathogen effectors target Arabidopsis Exo70B1, aiming to disrupt host secretion of defense-related molecules (30, 34, 35).

*Ralstonia solanacearum* is a destructive plant pathogen and the causative agent of bacterial wilt disease, affecting more than 250 plant species, including agronomically important crops (43–45). As in the case for many Gram-negative bacterial pathogens of both plants or animals, its Type III Secretion System (T3SS) is considered as a key virulence mechanism and is necessary for successful colonization of plant tissues by *R. solanacearum* (44, 46). T3SS is indispensable for the pathogen, since it serves for the injection of more than 70 Type III Effectors (T3Es) inside the host cell to promote colonization (44,47,48).

RipE1 is a T3E present in most of the so-far sequenced strains of the *R. solanacearum* species complex and seems to be highly conserved across different phylotypes (47,48). Recent data suggest the effector’s contribution to virulence and although it seems to trigger plant immunity in *Nicotiana benthamiana, Nicotiana tabacum* and *Arabidopsis thaliana*, other *R. solanacearum* T3Es are potentially able to suppress these responses (49–53). RipE1 has been shown to promote degradation of JAZ (jasmonate-ZIM-domain) repressors through its cysteine protease activity, inducing JA signaling and consequently suppressing SA responses, in accordance with the earlier characterized RipE1 homologue HopX1 (formerly AvrPphE) from *Pseudomonas syringae* (49, 54). In addition, cysteine protease activity of RipE1 is known to be required for triggering immune responses in both *N. benthamiana* and *A. thaliana* (51). Apart from their catalytic triad, HopX/AvrPphE homologues additionally share a conserved N-terminal domain, the so-called A-box, which is of unknown function but apparently necessary for effector function *in planta* (55). For RipE1, the A-box domain is also required for triggering immune responses in *N. benthamiana* (51).

In this work we aimed to uncover new targets of the conserved *R. solanacearum* T3E RipE1 and further investigate its contribution to the pathogen’s virulence. We discovered that RipE1 has multiple potential targets and among which, the *Arabidopsis thaliana* exocyst component Exo70B1 is a subcellular target of the particular effector *in planta*. We use AI-based effector structure and function predictions and demonstrate *in vitro* cleavage and *in planta* degradation of Exo70B1 by cysteine protease RipE1. In addition, we uncover that a previously characterized CC-NLR, named *Pseudomonas tomato race 1* (Ptr1), mediates RipE1 recognition in *Nicotiana benthamiana*. Overall, this study grants novel insights into discovering plant pathogen effectors’ distinctive targets and is the first report of a *R. solanacearum* effector targeting a component of the host secretion machinery. Finally, this target led to the revealing of the corresponding NLR able to activate cell death in non-host plants upon RipE1 recognition.

## Results and Discussion

### RipE1 intracellular function based on structure prediction

The AlphaFold, an artificial intelligence (AI), developed by DeepMind, protein structure prediction tool and it has been proposed as a highly accurate method to predict protein structures, compared to other well established methods (56). The 3D structure of RipE1 (47.31 kDa) was predicted using the AlphaFold Colab (**Fig. 1A**). The resulted structure confirmed that RipE1 has a cysteine protease folding. Cysteine proteases (thiol proteases) are enzymes with the ability of protein degradation by deprotonation of a thiol in the enzyme’s active site (57). Therefore, all enzymes in this category have a common catalytic triad or dyad involving a nucleophilic cysteine residue (57). The predicted RipE1 structure was compared to Protein Data Bank structures via the DALI online server (58). The highest similarity was with the *Legionella pneumophila* periplasmic protease LapG (59) (**Fig. 1C**). According to this comparison, the extending hydrophobic surface patch at the active site of RipE1 (**Fig. 1B**) may suggest a role as an interaction interface for substrate binding. Subsequently, the superimposition (**Fig. 1D**) of the domain of the active sites between the two proteins had a root-mean-square deviation (RMSD) of 1.544. The catalytic triad cysteine-histidine-aspartate (C-H-D) of RipE1, specifically C172-H203-D222 (**Fig. 1A**), appeared to be identical with the LapG C137-H172-D189 catalytic triad (**Fig. 1D**). RipE1 belongs to the HopX1/AvrPphE effector family, the members of which share a conserved N-terminal domain and a catalytic triad with predicted cysteine protease activity (54,55,60). RipE1 has been shown to possess cysteine protease activity *in vitro* and it seemingly depends on it to cleave JAZ repressors (49). Using AI-based structural predictions (**Fig. 1**), we further confirm that RipE1 is a cysteine protease. This further supports the findings of Sang et al (51), who reported that RipE1 loses its immunity-inducing attribute in *Nicotiana benthamiana* and *Arabidopsis thaliana* plants, when the cysteine residue belonging to its catalytic triad, is substituted.

**Figure 1.**
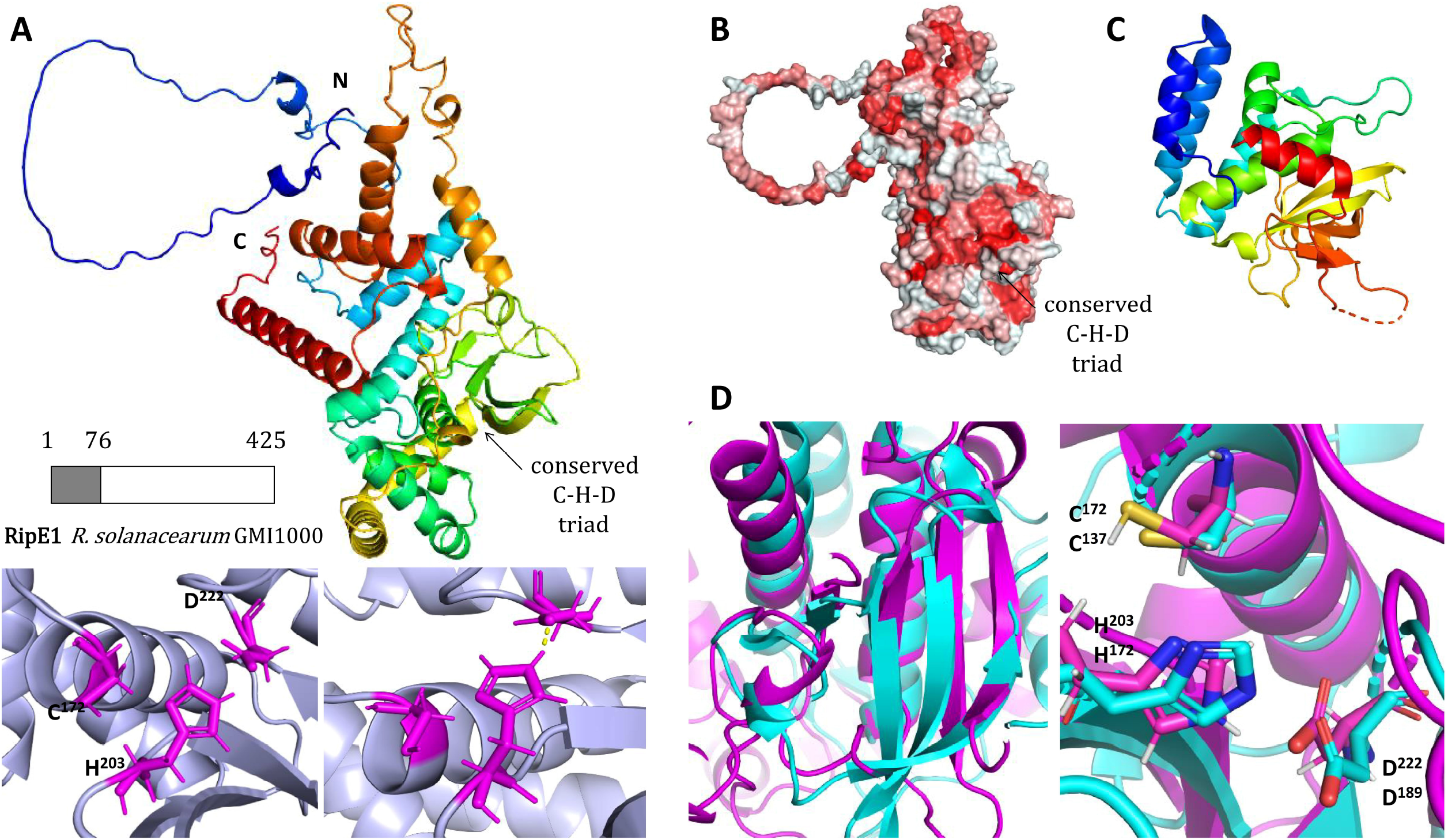
Presentation of *R. solanacearum* GMI1000 RipE1 3D structure prediction. **(A)** 3D structure prediction using AlphaFold Colab (N-terminal is shown as blue, C-terminal is shown as red). The first 76 residues are presented as unfolded. The conserved catalytic triad (Cys-His-Asp, C-H-D) and the hydrogen bond between H203 and D222is shown bottom left. **(B)** Surface hydrophobicity (Red: Hydrophobic surface). **(C)** Crystal structure of *L. pneumophila* LapG. **(D)** Superimposition of LapG (cyan) and RipE1 (purple) proteins active sites.

### RipE1 interacts with six ID protein domains and with Arabidopsis Exo70B1 in yeast

To identify new putative subcellular targets of RipE1, we performed a yeast two-hybrid (Y2H) screening between RipE1 and 22 eukaryotic protein domains (**Table S1**) that are fused to plant NLRs as Integrated Domains (NLR-IDs). We uncovered a total of 6 interactions (**Fig. 2A**) between RipE1 and NLR-IDs containing conserved characteristic domains of large plant protein families. RipE1 interacted with a TRX-like, a WRKY-like, the Exo70-like, and the CG-like NLR-IDs of our list, while weak interactions were also detected between RipE1 and the TFIIS-like, HSF-like and Ubox-like domains (**Fig. 2A, Table S1**). The expression of either RipE1 or any of the six NLR-IDs in yeast were not able to activate the reporter gene in the absence of an interactor, shortening the possibility of false-positive results (**Fig. S1A**). The conserved eukaryotic domains that interacted with RipE1 are commonly found on proteins that are tightly involved in regulation and facilitation of plant innate immunity processes (23,61–64).

**Figure 2.**
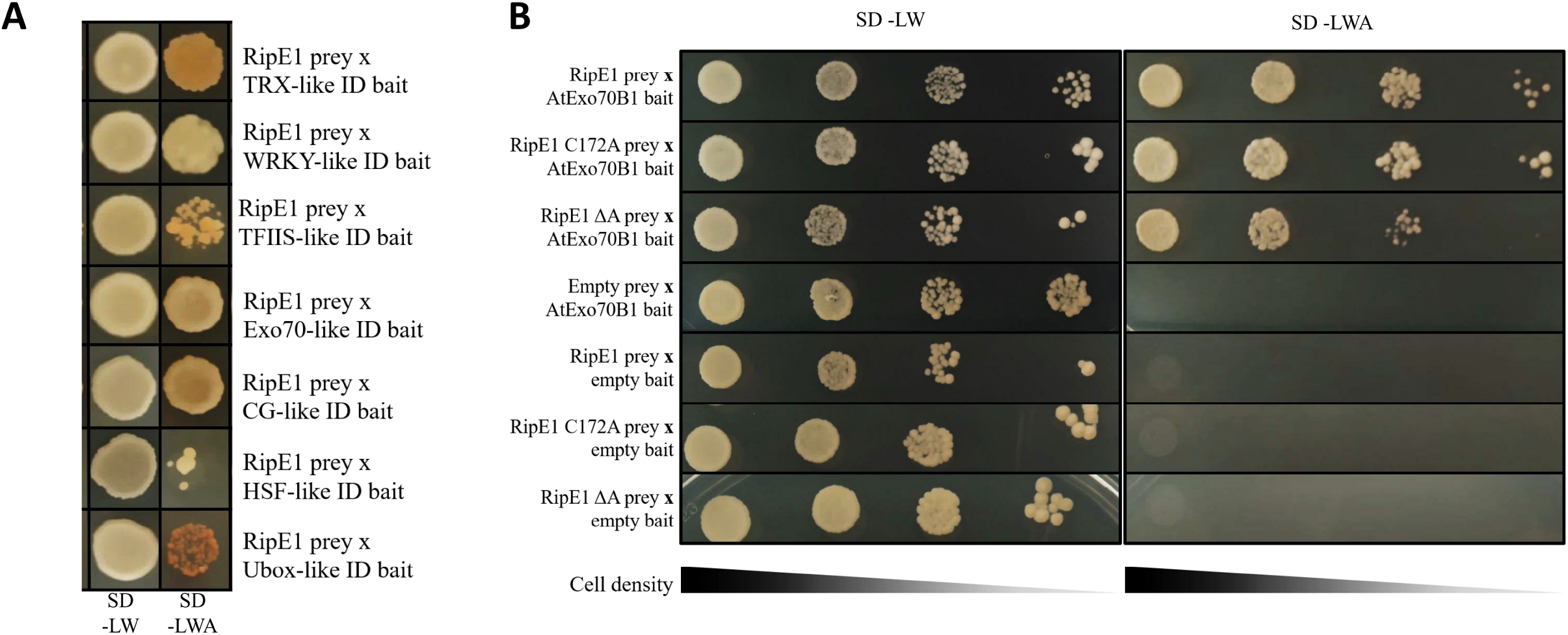
RipE1 interacts with Exo70-like ID and *At*Exo70B1 in yeast. **(A)** Interactions between RipE1 and six NLR-ID domains, uncovered by yeast two-hybrid screening (for more details see **Table S1**). Yeast cells carrying the indicated baits and preys were spotted on SDC-LW (plasmid selection) and SDC-LWA plates (interaction selection). Photographs were taken after 4 days of incubation. The red color in some of the colonies is a result of accumulation of a derivative of 5-aminoimidazole ribotide (87,88). **(B)** Interactions between wild-type RipE1, RipE1-C172A mutant, RipE1 ΔA mutant and *Arabidopsis thaliana* Exo70B1. Serial dilutions of cells carrying the indicated baits and preys were spotted on SDC-LW and SDC-LWA selective plates. Initial OD_600_ of cells was 1 unit and subsequent dilutions were made 0.1x, 0.01x and 0.001x. Photographs were taken after 4 days of incubation.

Pathogen effectors often interfere with multiple host pathways to promote virulence (4) and our results, along with the pre-existing literature, conjointly point out that RipE1 seems to be implicated in several T3E functions (49–51). Our findings additionally provide new putative RipE1 subcellular targets and we hope that they advance the discovery of more functions of this conserved *R. solanacearum* effector.

For the purposes of this study, we further investigated the interaction between RipE1 and Exo70-ID, aiming to uncover an effector target that belongs to the host secretory pathway. Based on our previously performed Blastp analysis using the Exo70-like ID as a query (34), we hypothesized that *At*Exo70B1, an Exo70-containing *Arabidopsis thaliana* protein and a component of the plant exocyst complex, is most likely to be targeted by RipE1. We proceeded by testing the interaction between Exo70B1 and RipE1 in yeast. As shown in **Figure 2B**, Exo70B1 not only interacted with RipE1, but also with two mutants of the effector, the RipE1-C172A, carrying a C172A substitution at the active site and the RipE1 ΔA lacking eight conserved amino acids (121-128) found in the N-termini of the HopX1 effector family, also known as the A-box (51,55). The two mutations were designed according to the structural predictions presented in **Figure 1**, but also due to their previously reported inability to induce plant immunity responses in model plants (51).

### RipE1 interacts with *At*Exo70B1 *in planta*

RipE1 has been shown to trigger a hypersensitive response in *Nicotiana benthamiana, Nicotiana tabacum* and induce defense responses in *Arabidopsis thaliana* (50,51). In this study, we screened various *Nicotiana* species, including various *Nicotiana sylvestris* ecotypes, to find a new plant model for our *in-planta* experiments. We discovered that RipE1 did not trigger an HR response when transiently expressed in leaves of the *Nicotiana sylvestris* ecotype NS2706 from our collection, via *Agrobacterium*-mediated infiltration (**Fig. S1B**) and thus, we used it as a novel tool and performed all our further experiments on this ecotype. To test the possible colocalization of RipE1 and Exo70B1 *in planta*, we transiently expressed both proteins fused to fluorescent tags, -YFP and - mCherry respectively, in *Nicotiana sylvestris* leaf tissue, observed at 48hpi via confocal microscopy. As shown in **Figure 3A**, RipE1 and Exo70B1 colocalized in the periphery of the plant cells, an observation supported by the *in-silico* colocalization analysis presented in **Figure 3B**. To further investigate the interaction between RipE1 and Exo70B1 *in planta*, we used Bimolecular Fluorescent Complementation (BiFC) assay, transiently co-expressing RipE1 tagged with C-terminal YFP (cCFP) and Exo70B1 tagged with N-terminal YFP (nVenus). YFP signal was detected in the cell periphery via confocal microscopy 48hpi, as shown in **Figure 3D**, indicating the association between the two proteins *in planta*. YFP signal was also detected with the RipE1 inactive mutant, RipE1-C172A, but not with the RipE1 ΔA mutant **(Fig. S2A**). Deletion of the A-box, had previously been shown to abolish cell death induction by RipE1 in *N. benthamiana* (51). This domain, although conserved in the HopX1/AvrPphE effector family, is of unknown function and has been earlier proposed as a domain that facilitates effector-target interactions (54,55). Our results show that deletion of this 8-amino acid domain does not alter RipE1 physical interaction capacity with Exo70B1 in *ex-vivo* assays (**Fig. 2B**) but it does abolish its ability to associate with Exo70B1 *in planta* (**Fig. S2A**). Interestingly, our data reveal an altered effector’s subcellular localization (**Fig. S1C**). Thus, its role in effector function remains to be elucidated. We additionally performed co-immunoprecipitation assays to validate RipE1-Exo70B1 interaction. Exo70B1-HF pull-down did precipitate RipE1-myc, after their *Agrobacterium*-mediated transient co-expression (**Fig. 3C**), advocating the association between the two proteins in the plant cells. These findings are derived from the Y2H screening presented and discussed above (**Fig. 2, Table S1**). After further investigation of the interaction between RipE1 and Exo70-ID, we established *Arabidopsis thaliana* exocyst component Exo70B1 as a subcellular interactor of RipE1, highlighting the benefits of employing NLR-ID domains as the means to uncover new potential targets of pathogen effectors.

**Figure 3.**
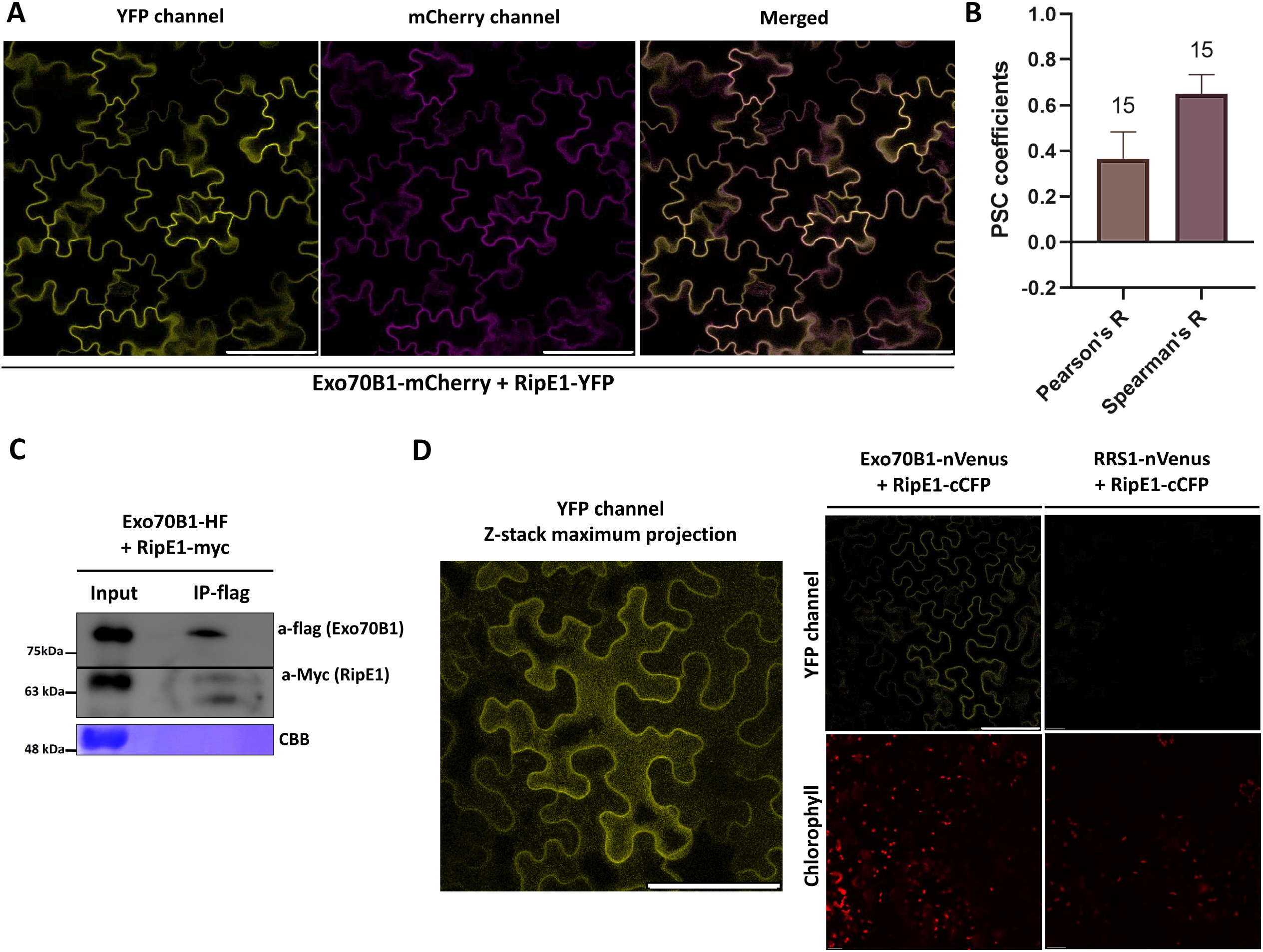
RipE1 interacts with *At*Exo70B1 *in planta*. **(A)** Colocalization between RipE1 and Exo70B1 *in planta*. YFP-tagged RipE1 and mCherry-tagged Exo70B1 were transiently expressed in *Nicotiana sylvestris* leaves and detected via confocal microscopy 48hpi. YFP and mCherry signals were overlapping at the cell periphery. **(B)** Colocalization analysis, supporting **Figure 1A**. 15 ROIs were analyzed using Coloc2 in ImageJ and the Pearson’s R and Spearman’s R parameters were plotted to visualize colocalization probability. **(C)** Co-immunoprecipitation of RipE1 and Exo70B1 from plant tissue. Myc-tagged RipE1 and HF (6xHis and 3xFLAG)-tagged Exo70B1 transiently expressed in *N. sylvestris* leaves and an IP-flag assay was performed 48hpi. RipE1-myc is 52 kDa and Exo70B1-HF is 77 kDa. **(D)** *In planta* validation of the RipE1/Exo70B1 interaction using a BiFC assay in *N. sylvestris* plants. RipE1 and Exo70B1 were fused at their C-termini with cCFP and nVenus epitope tags, respectively, and the YFP signal was detected via confocal microscopy 48hpi (middle). A maximum projection of the z-stack is also presented (left). As negative control, RipE1-cCFP was co-expressed with RRS1-nVenus and no YFP signal was observed (right). Bars = 80μm.

### RipE1 cleaves *At*Exo70B1 *in vitro*

To expand our understanding of the interaction between Exo70B1 and RipE1, we predicted and visualized the interaction *in silico* (**Fig. 4A**). Based on this, the Exo70B1 146-201 aa region appears to approach the catalytic triad of RipE1 cysteine protease. This observation, along with the interaction between the two proteins (**Fig. 2B, Fig. 3**), indicate that Exo70B1 is a substrate for RipE1. In order to test the proteolytic effect of RipE1 on Exo70B1, *in vitro* cleavage assays were performed with the purified proteins (**Fig.4B**). Purified 6xHis-SUMO3-Exo70B1 was incubated with RipE1-6xHis, with and without the addition of cysteine protease inhibitor leupeptin. Each protein of interest and the 6xHis-SUMO3 tag were all incubated separately, to serve as negative controls for the assay. The reactions were analyzed via SDS-PAGE, followed by anti-his-tag western blotting (**Fig. 4B**). RipE1 appeared to have autoproteolytic activity in all cases, including in the presence of leupeptin. On the other hand, a band representing cleaved 6xHis-SUMO3-Exo70B1 (see **Fig. 4B**, orange box) was detected only in absence of this inhibitor and in presence of RipE1 (**Fig. 4B**).

**Figure 4.**
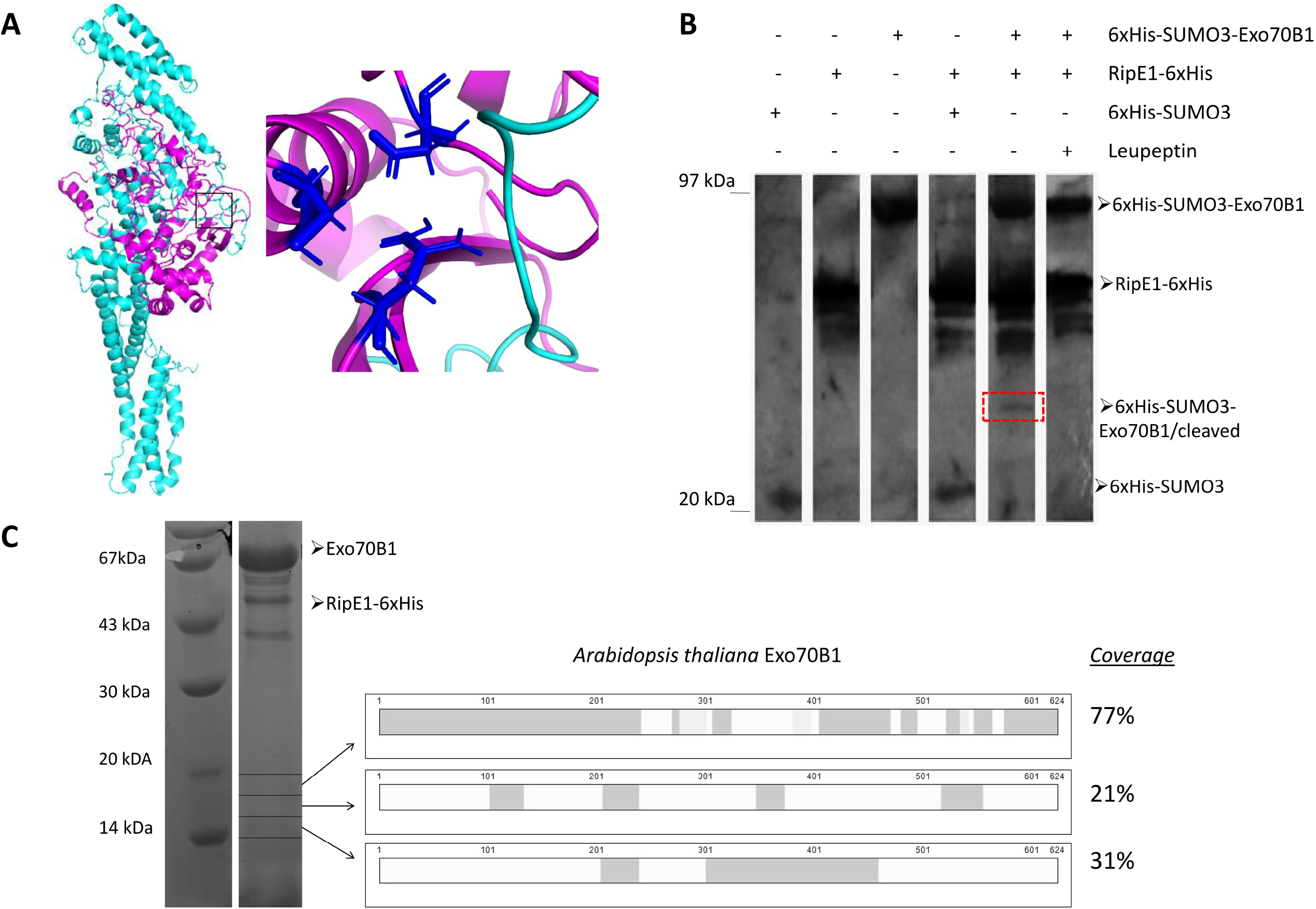
RipE1 cleaves Exo70B1 between its sequence *in vitro*. **(A)**. *In-silico* prediction of the interaction between Exo70B1 (cyan) and RipE1 (magenda). The boxed area appears magnified on the right and shows the Exo70B1 region (cyan) that approaches the catalytic triad of RipE1 (blue). **(B)** *In-vitro* cleavage of Exo70B1 by RipE1 is shown in this immunoblot. Reactions were analyzed via SDS-PAGE, followed by immunoblotting with His antibody. **(C)** Purified Exo70B1, after removal of 5xHis-SUMO3 tag, was incubated with RipE1-6xHis and the sample was analyzed via SDS-PAGE followed by Silver staining (left). Three selected SDS-PAGE zones, as indicated by arrows, were further analyzed by mass spectrometry. Grey regions on the right show the MS-identified Exo70B1 peptides that derived after cleavage by RipE1.

The *in-vitro* cleavage assay was repeated, after His:SUMO3 tag was removed from Exo70B1. The reaction was analyzed via SDS-PAGE, followed by Silver staining (**Fig. 4C**, left). Based on the cleaved 6xHis-SUMO3-Exo70B1 (**Fig. 4B**) band molecular weight and with the His-SUMO3 tag having been removed, three zones were selected to be further analyzed via mass spectrometry, marked by the arrows on **Figure 4C**. The analysis of the MS-derived data identified peptides of Exo70B1 present in these zones, as indicated by the grey areas (**Fig. 4C**, right). These findings confirmed that Exo70B1 is cleaved by RipE1 *in vitro*, between the particular sequence.

### RipE1 promotes the degradation of Exo70B1 *in planta*

To investigate the possible degradation of Exo70B1 by RipE1 *in planta*, we transiently expressed the proteins of interest in *N. sylvestris* tissue via *Agrobacterium*-mediated infiltration. Proteins were extracted and analyzed via SDS-PAGE and immunoblotting (**Fig. 5A**) and a total of three replications were used for *in-silico* quantification of proteins band intensity (**Fig. 5B**). In agreement with the observations regarding *in vitro* cleavage (**Fig. 4**), expression of RipE1 seemed to decrease Exo70B1 protein levels after their transient co-expression *in planta*, in a statistically significant manner (**Fig. 5B**). Compared with the cleavage of Exo70B1 by RipE1 *in vitro*, its *in-planta* degradation appeared more considerable. This complies with the fact that the *in vitro* experiments presented in **Figure 4** probably do not mirror the exact native conditions of the interaction between RipE1 and Exo70B1, while the experiments presented in **Figure 5** were conducted in plant tissue.

**Figure 5.**
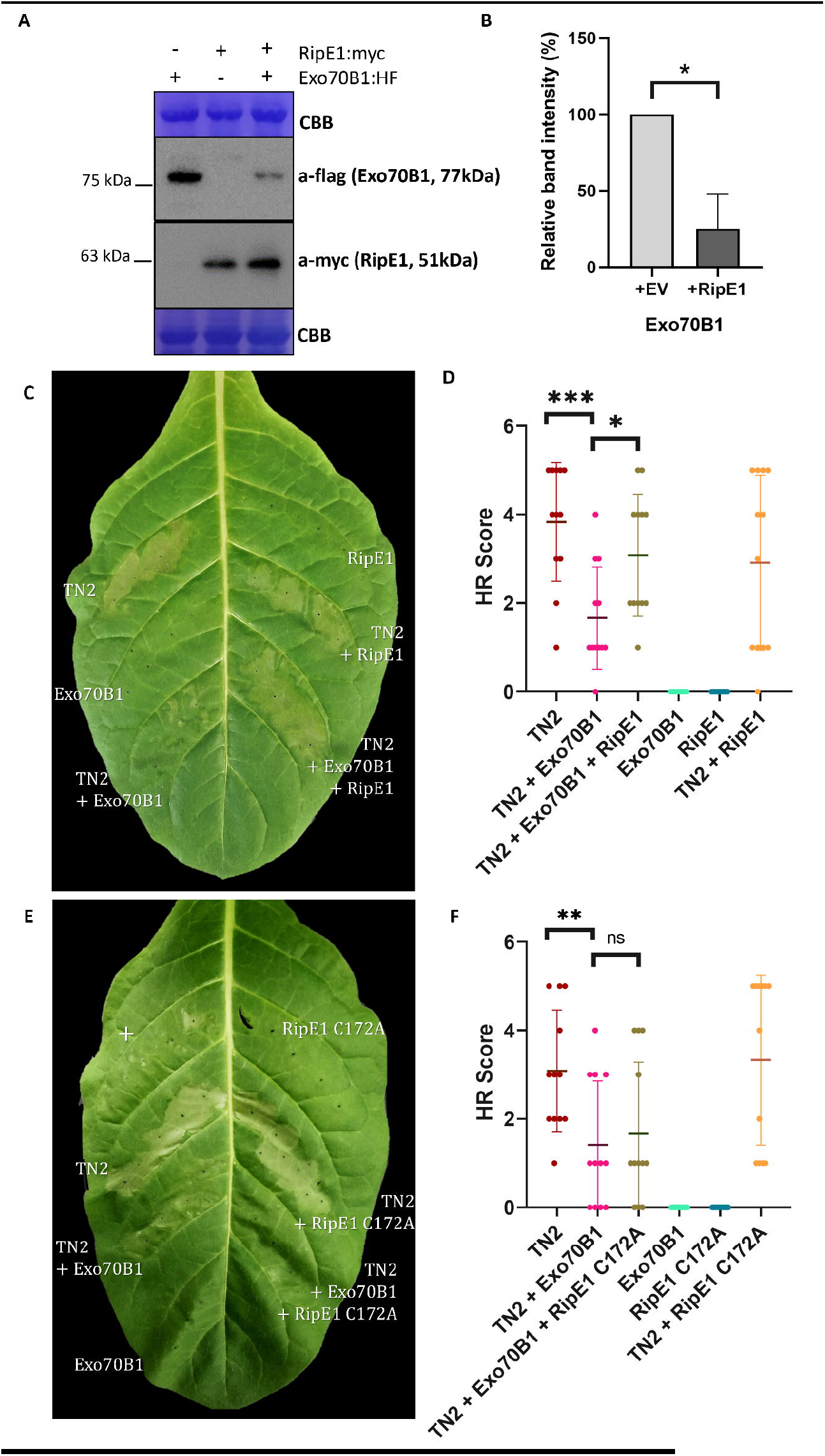
RipE1 promotes degradation of Exo70B1 *in planta*. **(A)** RipE1 expression promotes decrease in Exo70B1 protein levels in planta. These immunoblots support the statistical analysis presented in (B) and are one of the 3 replications performed for this analysis. RipE1 fused to Myc epitope and Exo70B1 fused to HF (6xHis and 3xFlag) epitope were transiently expressed in *Nicotiana sylvestris* leaves. Extracted proteins (48hpi) were analyzed via SDS-PAGE and immunoblotting to compare protein levels between the tissue samples expressing the single proteins (empty vector was used to balance agrobacteria density) and those co-expressing the interacting partners RipE1/Exo70B1. The presented analyzed samples have originated from the same leaf, to achieve comparability. Coomassie Brilliant Blue (CBB) staining images accompany each immunoblot and were used to normalize protein levels according to loaded protein quantity. **(B)** Statistical analysis of relative immunoblot band intensity indicating Exo70B1 protein levels extracted from plant tissue. This analysis was performed using 3 total experimental replicates, where total protein was extracted from *N. sylvestris* leaves, after *Agrobacterium*-mediated transient expression (48hpi) of Exo70B1 and Exo70B1 along with RipE1. Empty vector (EV) was used in the samples expressing only Exo70B1 to balance agrobacteria density. Compared relative band intensities (%) were submitted to t-test followed by Welch’s correction. *p value=0.0284 **(C)** *Nicotiana sylvestis* leaf representing one of the 12 replicates used for the statistical analysis of hypersensitive response (HR) assays presented in (D). The indicated proteins were transiently expressed via *Agrobacterium*-mediated infiltration at a total OD_600_ of 0.5. The photograph was captured 3dpi. **(D)** Statistical analysis of the 12 HR assays replicates testing wild type RipE1 ability to rescue TN2-induced cell death phenotype via Exo70B1 degradation. Compared phenotypes were submitted to t-test followed by Welch’s correction. ***p value=0.0003, *p value=0.0125. **(E)** *N. sylvestis* leaf representing one of the 12 replicates used for the statistical analysis of HR assays presented in (F). The indicated proteins were transiently expressed via *Agrobacterium*-mediated infiltration at a total OD_600_ of 0.5. The photograph was captured 3dpi. (+) on the upper left infiltrated area represents an additional cell-death inducing construct that was tested as a putative positive control (effector Chp7 fused to PR1 secretion peptide) and was not included in the statistical analysis of HR scores. **(F)** Statistical analysis of the 12 HR assays replicates testing mutated RipE1-C172A ability to rescue TN2-induced cell death phenotype via Exo70B1 degradation. Compared phenotypes were submitted to t-test followed by Welch’s correction. **p value=0.0085, ns: no significance p value>0.05.

There is strong evidence that Exo70B1 is guarded by the TN2 (TIR-NBS2) NLR in Arabidopsis in a CPK5 (Calcium-dependent Protein Kinase 5)-dependent manner and that Exo70B1 degradation induces NLR-mediated immunity responses (30, 39, 59). TN2 expression in *Nicotiana tabacum* triggers a strong cell death response, suppressed by co-expression with Exo70B1 and subsequently rescued by the *Pseudomonas syringae* effector AvrPtoB, which induces Exo70B1 degradation (30). Based on our findings so far, we hypothesized that Exo70B1 degradation by RipE1 would be able to induce TN2-dependent cell death response. We observed that TN2 also triggers a cell death response in *Nicotiana sylvestris* that was successfully suppressed by co-expression of Exo70B1 (**Fig. 5C-5F**). In accordance with our hypothesis, co-expression of TN2 and Exo70B1 with RipE1 did rescue TN2-mediated HR in *N. sylvestris* (**Fig. 5C-5D**), suggesting that RipE1 promotes Exo70B1 degradation *in planta*. When the RipE1 mutant C172A was used to perform similar assays, RipE1-C172A failed to rescue TN2-mediated HR, indicating that this rescue is dependent on the effector’s cysteine protease activity (**Fig. 5E-5F**). This NLR activation could possibly cause the C172-dependent induction of immune responses by RipE1 in Arabidopsis, as it has been reported by Sang et. al (51). In said study, the authors reported induction of immunity-related marker genes after RipE1 expression in transgenic Arabidopsis plants, but they did not observe activation of cell death (51). Interestingly, in our results, although RipE1 was able to activate TN2-dependent cell death when the receptor was ectopically expressed in *N. sylvestris* leaves (**Fig. 5E-5F**), transient expression of RipE1 in Arabidopsis Col-0 leaves failed to activate cell death (**Fig. S3**). To explain these observations, further knowledge is required regarding the regulation of TN2-activated hypersensitive response in Arabidopsis.

### RipE1 is recognized by *Nicotiana benthamiana* NLR Ptr1a, via its cysteine protease activity

Although cell death activation by RipE1 in *Nicotiana benthamiana* tissue has been previously reported, the NLR required for this activation has not yet been identified (51). RipE1-induced cell death in *N. benthamiana* seems to be independent of EDS1 (51) and NRG1 (**Fig. S2B**). Since both EDS1 and NRG1 have been linked to signaling downstream to TIR-NLRs (TNLs) activation, it is reasonable to hypothesize that the NLR responsible for RipE1 recognition in *N. benthamiana* is a non TIR-NLR and very likely it is a CC-NLR (CNL) (66). In addition, considering RipE1 ability to activate cell death or not in different *Nicotiana* species, a homolog of this ‘suspected’ NLR should not be present in *N. sylvestris* genome, but it should be found in *N. tabacum*. No such NLR has been associated with monitoring Exo70 proteins integrity in tobacco plants. However, it has been shown before that the CC-NLR, *Pseudomonas tomato race 1* (Ptr1) present in *N. benthamiana*, which harbors such characteristics, is associated with cleavage of RPM1-interacting protein 4 (RIN4) (67,68). Based on this and on various accumulating evidence regarding the association of RIN4 with Exo70B1 and other exocyst components (33,38,69,70), we investigated whether *Nb*Ptr1 is able to recognize RipE1. The CC-NLR, RPM1, has also been linked to the guarding of RIN4 against the *P. syringae* effectors AvrB and AvrRpm1 (71). However, an RPM1 homolog was not discovered in *N. benthamiana* and *N. tabacum* genomes in our genome mining analysis and this was also the case for a third RIN4-guarding NLR, the Arabidopsis *Resistance to Pseudomonas syringae 2* (RPS2), which guards RIN4 against AvrRpt2-mediated cleavage (72). We therefore hypothesized that the most suitable candidate NLR to test for RipE1-induced cell death in *N. benthamiana* was *Nb*Ptr1. To test our hypothesis, we transiently co-expressed *Nb*Ptr1 with RipE1 in *N. sylvestris* leaves via *Agrobacterium*-mediated infiltration (**Fig. 6A-6B**). Interestingly, the ectopic expression of *Nb*Ptr1 in *N. sylvestris* did not activate cell death when it was expressed without RIN4. Furthermore, the co-expression of *Nb*Ptr1 with the wild type RipE1 in *N. sylvestris*, activated a strong cell death phenotype, while such a phenotype was not observed when the inactive mutated RipE1-C172A was co-expressed with *Nb*Ptr1 (**Fig. 6A-6B**). These results indicate that *Nb*Ptr1 mediates recognition of RipE1, dependent on the effector’s cysteine protease activity, when ectopically expressed in *N. sylvestris*. To confirm RipE1 recognition by *Nb*Ptr1 in *N. benthamiana*, we used an *Agrobacterium*-mediated hairpin-based silencing system (73) to transiently silence *Nb*Ptr1 gene expression (**Fig. 6C**). RipE1 transient expression failed to activate cell death, when infiltrated in a *N. benthamiana* leaf area where an *Nb*Ptr1-targeting silencing hairpin had been priorly expressed (**Fig. 6C**). Overall, these findings suggest that *N. benthamiana* CC-NLR *Nb*Ptr1 is required for the activation of cell death by RipE1 cysteine protease activity.

**Figure 6.**
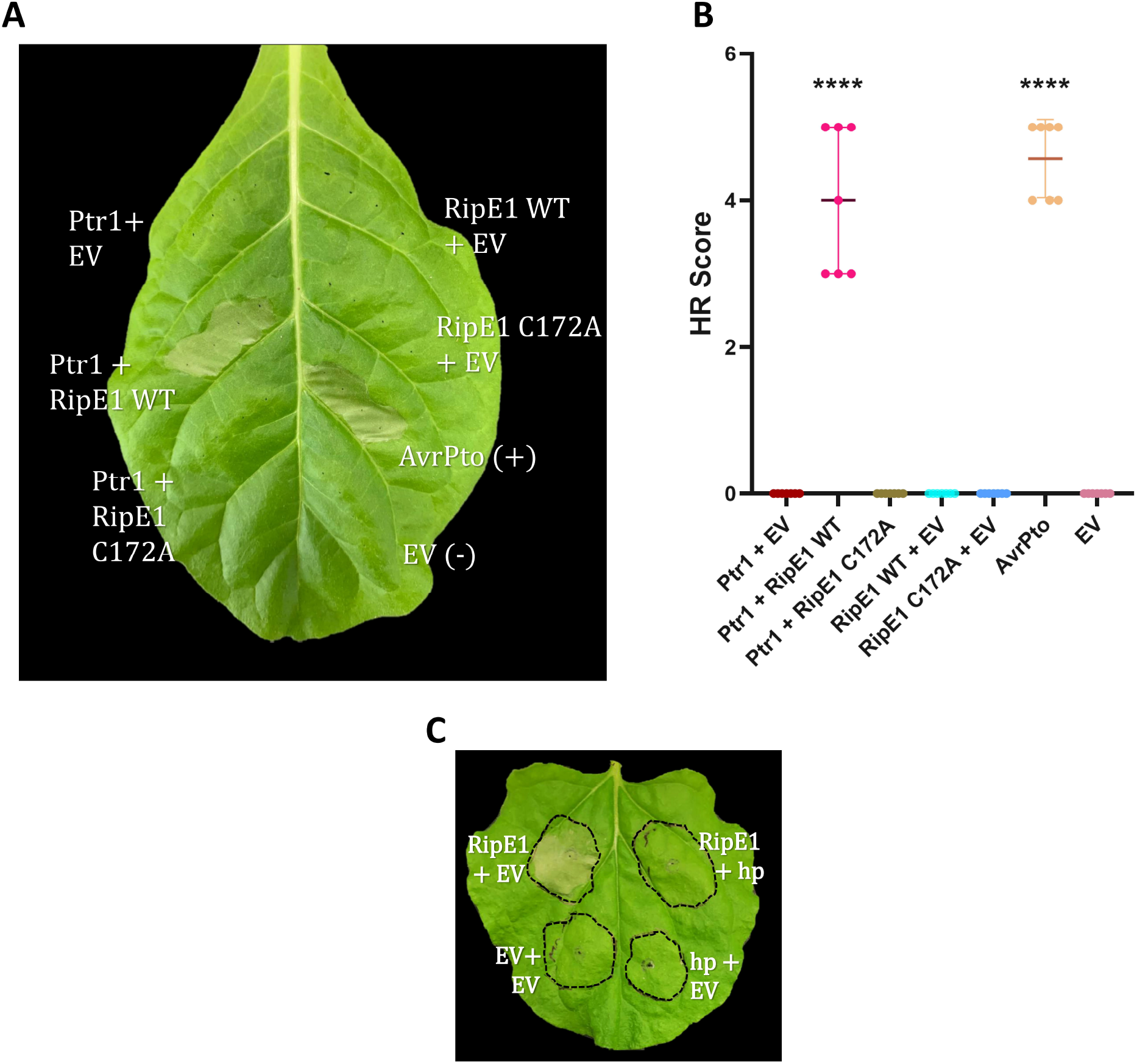
RipE1 is recognized by *N. benthamiana* NLR Ptr1 based on RipE1 protease activity. **(A)** *Nicotiana sylvestris* leaf representing one of the 7 replicates used for the statistical analysis of hypersensitive response (HR) assays presented in (B). The indicated proteins were transiently expressed via *Agrobacterium*-mediated infiltration at a total OD_600_ of 0.5. AvrPto transient expression was used as an HR-activating positive control. Empty vector (EV) was used as a negative control and also to balance total inoculum OD_600_.The photograph presented here was captured 2dpi. **(B)** Statistical analysis of the 7 HR assays replicates testing RipE1 ability to activate *Nb*Ptr1a-induced cell death phenotype via its cysteine protease activity. Compared phenotypes were submitted to t-test followed by Welch’s correction. ****p value<0.0001. **(C)** Silencing of *Nb*Ptr1 abolishes RipE1-induced cell death in *N. benthamiana*. This experiment was performed according to Brendolise et al., where agroinfiltrations are performed in two subsequent days. The first days, the two right patches were infiltrated with agrobacteria carrying the hairpin construct (hp), while the two left patches with agrobacteria carrying the corresponding empty vector (EV). The following day, the two upper patches were infiltrated with agrobacteria carrying a RipE1 construct, while the two bottom patches with agrobacteria carrying the corresponding empty vector. Photographs were taken 3 days after RipE1 infiltration.

### The guarded RIN4 is not a direct target of RipE1

*Nb*Ptr1 activation is associated with cleavage of RIN4 (67,68). Based on this, we decided to investigate whether RipE1 targets RIN4 in addition to Exo70B1 targeting. We tested the potential interaction between RipE1 and RIN4, initially *ex vivo* using a Y2H assay (**Fig. 7A**) and *in planta* via BiFC assay (**Fig. 7B**). RipE1 did not associate with RIN4 in yeast (**Fig. 7A)**. However, a quite weak YFP signal was observed in some cells when a BiFC assay was performed to test RipE1-RIN4 association *in planta* (**Fig. 7B**). The signal was stronger and detected in most cells for the RipE1-C172A/RIN4 association (**Fig. 7B**). To further investigate the possibility of RipE1 targeting RIN4, we attempted to visualize the two proteins *in silico* via AlphaFold, as it was done for RipE1/Exo70B1 complex (**Fig. 4A)**. The software could not predict a complex between RipE1 and RIN4 in a confident manner, thus the results of this analysis were inconclusive (data not shown). The data presented here overall suggest that RipE1 and RIN4 do not physically interact (**Fig.7A**), but they associate and are in close proximity *in planta* (**Fig. 7B**).

**Figure 7.**
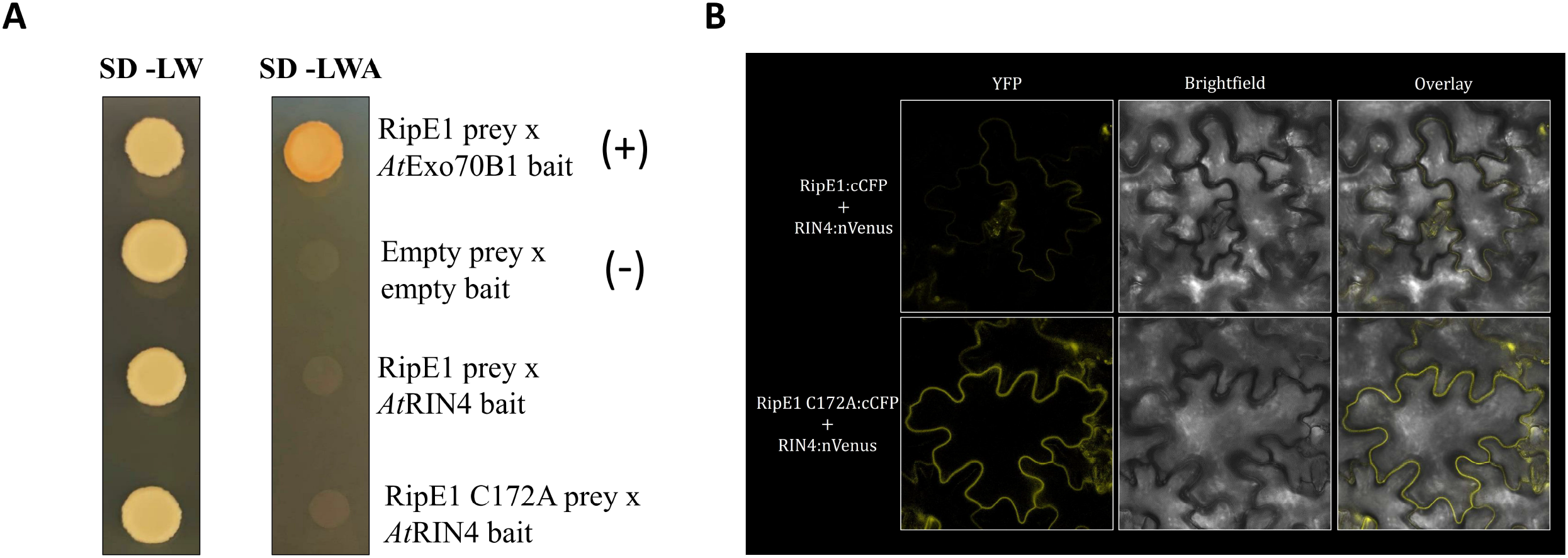
The guarded RIN4 is not a direct target of RipE1. **(A)** RipE1 and its derived mutants RipE1-C172A and RipE1 ΔA do not interact with Arabidopsis RIN4 in Y2H assay. RipE1 interaction with AtExo70B1 was used as a positive control. Yeast cells carrying the indicated baits and preys were spotted on SDC-LW (plasmid selection) and SDC-LWA plates (interaction selection). Photographs were taken after 4 days of incubation. **(B)** BiFC assays to test the association of RipE1 and its derived mutant RipE1-C172A with *At*RIN4. RipE1 proteins and RIN4 were transiently expressed in *N. sylvestris* leaf tissue via agroinfiltration, fused at their C-termini with cCFP and nVenus epitope tags, respectively. The YFP signal was detected via confocal microscopy 48hpi.

One plausible interpretation of these results is that RipE1 may indirectly compromise RIN4 via Exo70B1 degradation. In 2017, Sabol et al reported that upon AvrRpt2 delivery, both RIN4 fragments and Exo70B1 are released from the cell PM to the cytoplasm (38). Although these data support the hypothesis that RIN4 integrity is important for Exo70B1 localization to the PM, what happens to RIN4 localization in the case of Exo70B1 degradation has not yet been investigated. In addition, while RPM1 ectopic overexpression in tobacco results in cell death, unless RIN4 is also over-expressed (74,75), our results show that *Nb*Ptr1 ectopic over-expression in *N. sylvestris* does not elicit cell death (**Fig. 6A**). Considering this, *Nb*Ptr1 may monitor RIN4 without a physically association with it, leaving open the possibility that other RIN4-interacting proteins -like Exo70B1-are implicated in this process. Nevertheless, further investigation is required in order to decipher the activation of *Nb*Ptr1 by RipE1 and to determine if Exo70B1 degradation is involved in this mechanism. A proposed model describing a molecular mechanism, that collectively integrates the findings of this study, is illustrated in **Figure 8**.

**Figure 8.**
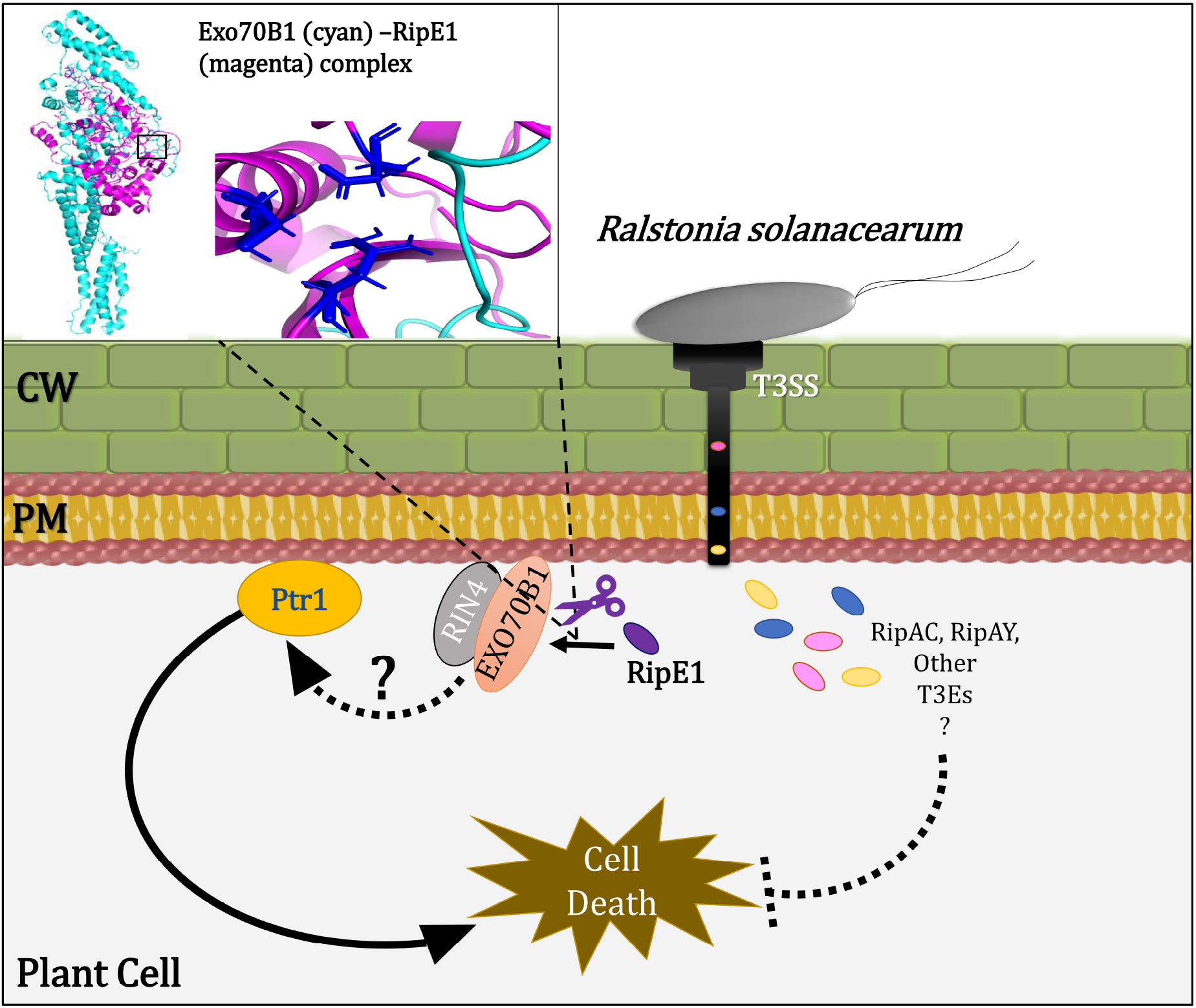
Schematical Abstract. *Ralstonia* infection, RipE1 is delivered into the cytoplasm among other effectors. Its cysteine protease activity is perceived by Ptr1, activating cell death. Proteolytic cleavage of Exo70B1 by RipE1 could lead to Ptr1-induced cell death due to Exo70B1/RIN4 interaction at the PM. Other T3Es secreted by *R. solanacearum*, like RipAC (53) or RipAY (51), can suppress RipE1-triggered cell death.

## Conclusion

Overall, the evidence presented here heavily supports the fact that RipE1 targets and cleaves the host cell’s exocyst component Exo70B1, possibly adding the manipulation of the host secretion machinery by RipE1 to the pathogen’s arsenal of virulence means. However, this plant protein is also involved in other eukaryotic pathways and given the displayed embroilment of Exo70B1 in plant autophagy (40), whether cleavage by RipE1 aims to interfere with vesicle trafficking or other host cell processes needs to be elucidated. In addition, our data uncovered an existing plant NLR of *N. benthamiana* that can recognize RipE1, thus contributing to pursuing resistance against *R. solanacearum*.

## Materials and Methods

### *In silico* protein structure prediction and analysis

The 3D structure of the full length RipE1 protein was predicted using the AlphaFold Colab (76,77). Subsequently, the predicted structure was compared with those in the Protein Data Bank via the Dali (58) online server. The visualization of the protein structures was performed by PyMOL (78).

### Plasmid construction

Genes of interest were amplified using Phusion® High-Fidelity DNA Polymerase (NEB) and PCR products were purified with the Nucleospin® Gel and PCR Clean-up kit MACHEREY-NAGEL). All constructs used in this study were generated via Golden Gate (GG) Cloning, using the following plasmids as backbone vectors: pGADT7:RFP, pGBKT7:RFP for yeast and pICH86988, pICH86966 for plant assays. RipE1 was amplified from *Ralstonia solanacearum* genomic DNA. *R. solanacearum* GMI1000 was kindly donated by Prof. Nemo Peeters (INRA-CNRS) and gDNA was extracted using the CTAB method (79). RipE1 mutations were designed according to structural predictions (**Figure 1**) and recent studies (49,51) and were generated through PCR followed by GG Cloning, using the primers listed in **Table S2**. Exo70B1 and NLR-ID containing constructs were generated as described earlier in a recent study from our lab (34). pMDC:spC7HPB for *in planta* transient expression of Chp7 was a kind donation of Prof. Jane Glazebrook (80). Ptr1a was amplified from *Nicotiana benthamiana* cDNA and was cloned in GG-compatible vectors using the primers listed in **Table S2**. All constructs generated in this study were verified by Sanger sequencing.

### Bacterial strains and growth conditions

*Escherichia coli* (Stellar or DH10b strains) were cultured in LB medium (1% tryptone, 0.5% yeast extract, 1% sodium chloride for liquid cultures and additionally 1,5% agar for plates) containing appropriate antibiotics: ampicillin 100μg/ml or kanamycin 50μg/ml. *E. coli* cells were grown either on LB plates or in liquid LB medium at 37°C for 16h. For liquid cultures, shaking was applied at approximately 200 rpm. *Agrobacterium tumefaciens* (AgL-1, C58C1 or GV3101 strains) were also cultured in LB medium containing appropriate antibiotics: rifampicin 50μg/ml and ampicillin 100μg/ml, kanamycin 50μg/ml or gentamycin 10μg/ml. *Ralstonia solanacearum* strain GMI1000 was cultured in NB/NA medium (0.5% tryptone, 0.3% yeast extract, 0.5% sodium chloride for liquid cultures and additionally 1,5% agar for plates) and grown at 28° for 2 days. For liquid cultures, shaking was applied at approximately 200 rpm.

### Y2H assay

All plant proteins were cloned as baits (in the pGBKT7-RFP vector) and bacterial effectors were cloned as preys (in the pGADT7-RFP vector), or as otherwise stated. Different pairs of constructs were co-transformed into *Saccharomyces cerevisiae* strain PJ694a (81), which carries the following auxotrophic traits: gal4Δ, gal80Δ, trp1-901 leu2-3 ura3-52 his3-200, *LYS2::GAL1-HIS3, GAL2-ADE2, met2::GAL7-lacZ*. At least 500ng from each plasmid (bait and prey) was used to transform *S. cerevisiae* PJ694a competent cells, freshly prepared with the lithium acetate method (82). Transformed cells were selected on SD–Leu–Trp medium (SD– LW) agar plates. Two single clones were transferred from the SD–LW medium and were re-cultured on SD–LW (for verification of growth) and SD–Leu–Trp–Ade (–LWA, auxotrophic selection) medium. For the dilutions, a single clone was transferred to liquid SD–LW medium, grown until saturation and diluted as stated in the figures, after OD_600_ measurements. Yeast cells in this study were cultured in YPDA rich medium or SD-LW and SD-LWA minimal media at 28-30°C.

### Plant material

The *Nicotiana sylvestris* plants from various ecotypes that were used in our study were grown in greenhouse conditions. For *in planta* transient expression via *Agrobacterium*-mediated infiltration, 4-week old *N. benthamiana*, 4-week old *A. thaliana* and 5-week-old *N. sylvestris* NS2706 plants were used. For agroinfiltration of *Nicotiana* plants, Agrobacteria carrying indicated constructs were grown in liquid cultures until saturation and cell pellets were diluted in 10mM MgCl_2_ and 10mM MES. Agrobacteria carrying each construct were then used at an OD_600_ score of 0.5 to infiltrate *N. sylvestris* leaf areas, while agrobacteria carrying empty vector were used to equalize the total OD_600_ score in all inoculums of each experiment. Agroinfiltration in Arabidopsis plants was performed as previously described (34,83).

### Confocal microscopy

All confocal images were captured in a Leica SP8 laser scanning confocal microscope. All leaf tissue samples were imaged with a 40x water objective, after *Agrobacterium*-mediated transient expression at 48-72hpi. The confocal images were processed with ImageJ. For colocalization, all indicated proteins were fused to YFP or mCherry fluorescent epitopes. YFP was excited using a 514-nm argon laser and emission was detected at 520–550 nm. mCherry was excited using a 561-nm laser and emission was detected at 593–628nm. The colocalization analyses were performed with PSC using the Coloc2 addon in Fiji (84,85). The threshold regression used was Costes, PSF 100 and Costes randomizations 10 (86). A total of 15 regions of interest (ROIs) were chosen for the PSC analysis. For BiFC, all indicated peoreins were fused to C-terminal nVENUS or cCFP epitope tags and subsequently, YFP signal was detected. Chlorophyll was detected between 653 and 676 nm and excited at 561 nm. Leica software LASX was used to add scale bars to the images and to display the z-stack maximum projection.

### Protein extraction from plant leaf tissue

Agroinfiltrated leaf tissue was ground at 48hpi in liquid nitrogen and proteins were extracted in GTEN buffer (10% glycerol, 25-mM Tris–Cl pH 7.5, 1-mM ethylenediaminetetraacetate, 150-mM NaCl), supplemented with fresh 5-mM DTT, protease inhibitor cocktail (Calbiochem) and 0.15% v/v NP-40, applying vigorous shaking. After a 10-minute centrifugation at high speed (14,000g) at 4°C, the supernatant was collected in a clean microcentrifuge tube and was used for further protein analysis (co-immunoprecipitation, SDS-PAGE and immunoblotting).

### Co-Immunoprecipitation

Protein extract from leaf tissue was incubated for 2h at 4°C with an anti-flag mouse antibody at 1:4,000 dilution (Sigma, F1804), followed by a 1h-incubation at 4°C with PureProteomeProtein A/G Magnetic Beads Mix (Millipore). Beads were washed 3 times with GTEN buffer, before proceeding to SDS-PAGE and immunoblotting.

### SDS PAGE and Immunoblotting

Protein samples from plant leaves were prepared for SDS-PAGE electrophoresis by adding 1x SDS loading dye and boiling the samples for 5 min. The samples were then separated via 8% or 12% SDS–PAGE (stated in images captions), electroblotted to a PVDF membrane (Millipore), then blocked for 1h with 5% w/v milk in Tris-buffered saline (TBS; 150-mM NaCl, 50-mM Tris–HCl, pH 7.6) and incubated for 1h with primary antibodies (1:4,000 dilution, anti-flag mouse -Sigma #F1804 or anti-myc mouse - Cell Signaling #2276). Membranes were then incubated for 1h with a secondary antibody (anti-mouse HRP conjugate at 1:10,000 dilution -Cell Signaling) and chemiluminescent substrate ultra (Millipore) was applied for 5 min before camera detection in Sapphire Biomolecular Imager (Azure Biosystems). For protein band intensity analysis, gel lanes (both CBB staining and immunoblot) from 3 separate experimental replications were selected and analyzed in FIJI v. 1.49 software (https://imagej.nih.gov/). Band intensity measurements were then submitted t-test followed by Welch’s correction on GraphPad Prism version 8.0.0 for Windows, GraphPad Software, San Diego, California USA, (www.graphpad.com). Differences with p value <0.05 were considered statistically significant (*), while multiple asterisks were used to indicate profoundly low p value (as stated in the corresponding figure legends).

### Protein production and purification

RipE1 gene was cloned and inserted into the pET26b (ColE1 plasmids) vector carrying a C-terminal 6-His tag and transformed into the *E. coli* strain BL21. A sufficient amount of soluble protein was obtained after induction using the following conditions: Cells were grown in LB medium containing 30μg/ml kanamycin until an OD_600_ value of 0.6-0.8 was reached. The culture was induced with 0.5 mM IPTG and left to grow at 20°C overnight. The cell paste was re-suspended in 100 ml lysis buffer containing 25 mM Tris pH 8.0, 300 mM NaCl, 5 mM imidazole and 15 mM β-mercaptoethanol, and homogenized. After adding protease inhibitors (20 μg/ml leupeptin, 1 mM PMSF and 150 μg/ml benzamidine), the cells were disrupted with sonication in ice, 10 sonication cycles of 30 s each, with cooling intervals of 30 s. The precipitate was subsequently removed by centrifugation at 12,000 rpm at 4°C for 45 min. Purification was carried out using His-tag affinity chromatography at 4°C with a 8 ml Ni-NTA Qiagen column pre-equilibrated in lysis buffer and initially washed stepwise with 10, 20 and 30 mM imidazole. With a subsequent increase in imidazole concentration the protein started eluting at 100 mM imidazole. Fractions containing the protein were dialyzed against the storage buffer containing 25 mM Tris pH 8.0, 100 mM NaCl and 10 mM β-mercaptoethanol. In case of EXO70B1, the full-length protein was cloned and inserted into the LIC 1.10 vector carrying a N-terminal 6-His-SUMO3 tag, and transformed into the *E. coli* strain Rosetta™(DE3). The LIC (ligation independent cloning) 1.10 vector carrying the tag was kindly donated by the Netherlands Cancer Institute (NKI). The same protocol was followed and the fractions containing the protein were dialyzed against the storage buffer (25 mM Tris pH 8.0, 100 mM NaCl, 10 mM β-mercaptoethanol and 0.5 nM SENP-2 protease). Subsequently, a reverse purification protocol was carried out using His-tag affinity chromatography. The elution fractions containing the proteins, were polled and concentrated to a final volume of 2 ml. Size exclusion chromatography was performed in 20°C ÄKTA purifier system (Amersham) and a prepacked Hi-Prep 16/60 Sephacryl S-200 high-resolution column (GE Healthcare). Flow rate was 0.5 ml/min, and elution was monitored at 280 nm. Using a 2ml-loop the proteins were loaded, and the 2ml fractions were collected and analyzed via a 10% SDS-polyacrylamide gel.

### *In vitro* cleavage assay

Purified *A. thaliana* Exo70B1 was incubated with purified RipE1 in reaction buffer (25 mM Tris pH 8.0, 100 mM NaCl), for 3 h at room temperature. For the inhibitory assay, RipE1 was incubated with 100 μM leupeptin prior the cleavage assay, at room temperature for 1 h before adding the substrate. After the indicated times, the reaction was stopped by the addition of SDS sample buffer, and the samples were subjected to SDS-polyacrylamide gel electrophoresis and analyzed by Western blotting.

### NanoLC-MS/MS Analysis

The nanoLC-MS/MS analysis was performed on an EASY-nLC II system (Thermo Scientific) coupled with an LTQ-Orbitrap XL ETD (Thermo Scientific) through a nanoES ion source (Thermo). Data were acquired with Xcalibur software (LTQ Tune 2.5.5 sp1, Thermo Scientific). Prior to the analysis, the mass spectrometer was calibrated with a standard ESI positive ion calibration solution of caffeine (Sigma), L-methionyl-arginyol-phenylalanylalanine acetate H2O (MRFA, Research Plus, Barnegat, NJ), and perfluoroalkyl triazine (Ultramark 1621, Alfa Aesar, Ward Hill, MA). Samples were reconstituted in 0.5% formic acid, and the tryptic peptide mixture was separated on a reversed-phase column (Reprosil Pur C18 AQ, particle size = 3 μm, pore size = 120 □ (Dr. Maisch, AnaLab, Athens, Greece), fused silica emitters 100 mm long with a 75 μm internal diameter (New Objective) packed in-house using a pressurized (35 to 40 bars of helium) packing bomb. The nanoLC flow rate was 300 nl min−1. The LC mobile phase consisted of 0.5% formic acid in water (A) and 0.5% formic acid in acetonitrile (B). A multi-step gradient was employed, from 5% to 30% B in 120 min, to 90% B in 10 min. After the gradient had been held at 90% B for 5 min, the mobile phase was re-equilibrated at initial gradient conditions. The MS was operated with a spray voltage of 2300 V, a capillary voltage of 35 V, a tube lens voltage of 140 V, and a capillary temperature of 180 °C. A survey scan was acquired in the range of m/z 400–1800 with an AGC MS target value of 106 (resolving power of 60,000 at m/z 400). The 10 most intense precursor ions from each MS scan were subjected to collision-induced dissociation in the ion trap.

### Data Analysis of MS/MS-derived Data

The MS raw data were loaded in Proteome Discoverer 2.2.0 (Thermo Scientific) and run using Mascot 2.3.02 (Matrix Science, London, UK) and Sequest HT (Thermo Scientific) search algorithms against the EXO70B1 Sequence. A list of common contaminants was included in the database. For protein identification, the following search parameters were used: precursor error tolerance = 10 ppm, fragment ion tolerance = 0.8 Da, trypsin full specificity, maximum number of missed cleavages = 3, and cysteine alkylation as a fixed modification, Oxidation of Methionine, Acetylation of N-term, Phosphorylation (The node ptmRS was used) of Serine, Threonine and Tyrosine as Variable Modifications. To calculate the protein false discovery rate, a decoy database search was performed simultaneously with strict criteria set to 0.01 and relaxed criteria to 0.05.

### Statistical analysis and plotting of Hypersensitive Response (HR) assay results

Statistical analysis and plotting were performed using GraphPad Prism version 8.0.0 for Windows, GraphPad Software, San Diego, California USA (www.graphpad.com). For HR assays, a score 0-5 was assigned to each observed phenotype, representing cell death intensity. HR scores from 12 replicates were submitted to t-test followed by Welch’s correction. Differences with p value <0.05 were considered statistically significant (*), while multiple asterisks were used to indicate profoundly low p value (as stated in the corresponding figure legends).

### *Agrobacterium*-mediated hairpin-based silencing

Silencing of *Nb*Ptr1 was performed via an *Agrobacterium*-mediated hairpin system, according to Brendolise et al. (73), where agroinfiltrations were performed in two subsequent days. The first day, the two right patches were infiltrated with agrobacteria carrying the hairpin construct (hp), while the two left patches with agrobacteria carrying the corresponding empty vector (EV). The following day, the two upper patches were infiltrated with agrobacteria carrying the effector construct, while the two bottom patches with agrobacteria carrying the corresponding empty vector. All constructs were infiltrated at OD_600_ of 0.5. Photographs were taken 3 days after effector infiltration.

## Acknowledgements

The authors want to thank Prof. Nemo Peeters for his kind donation of the *R. solanacearum* GMI1000 strain and Prof. Dimitris Goumas for his guidance with culturing and handling the bacteria; Prof. Despina Alexandraki for her generous donation of the yeast strain PJ69; Mr Marc Youles and the Synthetic Biology Group of TSL for their kind donation of Golden-Gate-Cloning compatible vectors for Y2H assays; Mrs Dina Kotsifaki for her valuable help during protein purification; Mr Nikos Kountourakis for his generous help with analysis of MS/MS-derived data; Dr. Patrick Celie for his kind donation of the LIC 1.10 vector; Dr. Cyril Brendolise for his generous donation of the *Nb*Ptr1 hairpin construct and lastly postdoctoral fellow Dr Glykeria Mermigka for her advice and fruitful discussion throughout the experiments.

## Author contributions

P.F.S. designed the research and supervised the project; D.T. and S.M. carried out the experiments; K.K. purified recombinant proteins and performed in vitro protein assays and performed the AlphaFold analysis; V.A.M. supervised the confocal microscopy experiments. M.K., V.A.M., contributed experimental materials. P.F.S., D.T. and K.K. wrote the manuscript. P.F.S. edited the manuscript. All authors have read and approved the manuscript.

## Supplementary Tables and Figures legends

**Table S1. NLR-IDs that were tested for interaction with RipE1**. NLR-ID domains originating from both dicot (*Arabidopsis thaliana, Brassica napus*) and monocot (*Oryza sativa, Hordeum vulgare*) plant genomes were used in a yeast two-hybrid screening to test their possible interaction with effector RipE1. Each putative interaction was tested at least 3 times.

(-) no interaction was detected

(+) a relatively weak interaction was detected, *not* detectable in all replications

(++) a relatively weak interaction was detected, detectable in all replications

(+++) a relatively strong interaction was detected, detectable in all replications

**Table S2. List of primers used in this study**. The primers listed here were designed to clone effector wild type RipE1 and its derived mutants RipE1-C172A and RipE1 ΔA, as well as the *Nb*Ptr1 gene, in Golden Gate-compatible vectors for expression in yeast (yeast two-hybrid assay, Y2H) and *Agrobacterium*-mediated transient expression in plant tissue.

**Figure S1. Representative images supplementing main Figures 2 and 3. (A)** Neither RipE1 or any of the NLR-IDs were able to induce expression of the reporter gene in a yeast two-hybrid assay, in the absence of an interacting protein. Empty bait and prey vectors, respectively, were used to test possible auto-activities. The results support the interactions in yeast, that are presented in **Figure 2. (B)** Wild type RipE1 or mutants C172A and ΔA do not induce a hypersensitive response (HR) in *Nicotiana sylvestris* selected ecotype NS2706. Effector Chp7 –fused to PR1 secretion peptide for apoplastic secretion-was used as a positive HR-inducing control and GUS reporter gene was used as a negative control. This photograph was taken 4dpi. This experiment was repeated at least 6 times with the same exact results. **(C)** Leaf discs from **(B)** were observed via confocal microscopy to confirm expression of proteins of interest that did not induce HR. All indicated proteins were c-terminally fused to YFP fluorescent epitope. These images were captured 5dpi. **(B) and (C)** collectively support the further use of *Nicotiana sylvestris* NS2706 as a model for our *in-planta* experiments.

**Figure S2. (A)** Investigation of the RipE1-C172A/Exo70B1 and RipE1 ΔA/Exo70B1 association using a BiFC assay in *Nicotiana sylvestris* plants. RipE1 mutated proteins and Exo70B1 were fused at their C-termini with cCFP and nVenus epitope tags, respectively, and the YFP signal was detected via confocal microscopy 48hpi. YFP signal was detected when RipE1-C172A-cCFP was co-expressed with Exo70B1-nVenus, while it was not detected upon co-expression of RipE1 ΔA-cCFP/Exo70B1-nVenus. **(B)** RipE1-induced cell death in *N. benthamiana* is independent of NRG1. Wild type RipE1 and the mutants RipE1-C172A and RipE1 ΔA were transiently expressed in 4-week-old *N. benthamiana nrg1* plants via agroinfiltration. The experiment was performed 3 times with identical results. Photograph was taken 3dpi.

**Figure S3. RipE1 does not activate cell death after transient expression in Arabidopsis leaves. (A)** *Arabidopsis thaliana* Col-0 representative leaf, after transient expression of effector RipE1. The dotted circle marks the infiltrated area. RipE1-myc was transiently expressed via *Agrobacterium*-mediated infiltration at a total OD_600_ of 0.5. Photographs were taken 7dpi. **(B)** RipE1-myc expression in (A) was confirmed via total protein extraction of both infiltrated (+) and non-infiltrated (-) areas and subsequent SDS-PAGE and western blot analysis.

