## Supplemental Table S2 for "The core effector RipE1 of *Ralstonia solanacearum* interacts with and cleaves Exo70B1 and is recognized by the Ptr1 immune receptor"

| Primer Name | Primer Sequence | Purpose of use |
| --- | --- | --- |
| ripE1-FW | aaGGTCTCaAATGCCGCCCGTCCTGCC | Cloning RipE1 |
| ripE1-REV | aaGGTCTCaAAGCTCAGCTTTCCGTGGCGGG | Cloning RipE1 for Y2H |
| tag_ripE1-REV | aaGGTCTCaCGAAggGCTTTCCGTGGCGGGc | Cloning RipE1 for C-terminal tagging (lacking a stop codon) |
| ripE1-C172A-REV1 | aaGGTCTCatccccgcccctgccaccattg | Cloning RipE1 with a C172A amino-acid substitution |
| ripE1-C172A-FW2 | aaGGTCTCagggaacGCcggcgaacacg | Cloning RipE1 with a C172A amino-acid substitution |
| ripE1-DelA-REV1 | aaGGTCTCagtgcgtcagtatccggcgggtt | Cloning RipE1 with an 8 amino-acid (121-128) deletion |
| ripE1-DelA-FW2 | aaGGTCTCagcacatcgacgccacccatggatt | Cloning RipE1 with an 8 amino-acid (121-128) deletion |
| NbPTR1A-FW1 | ttGGTCTCtAATGGCAGAATTTTTCTTGTTCA | Cloning NbPtr1a gene part 1 |
| NbPTR1A-REV11 | ttGGTCTCtACAGCCAAAGGCACTCCTC | Cloning NbPtr1a gene part 1 |
| NbPTR1A-FW2 | ttGGTCTCtCTGTGAAAACCTTGGGAAGGT | Cloning NbPtr1a gene part 2 |
| NbPTR1A-REV2tag | ttGGTCTCtCGAAggGCATCCTCCACTTAGCAATGG | Cloning NbPtr1a gene part 2 for C-terminal tagging (lacking a stop codon) |

**Suppl. Table S2. List of primers used in this study.** The primers listed here were designed to clone effector wild type RipE1 and its derived mutants RipE1 C172A and RipE1 ΔA, in Golden Gate-compatible vectors for expression in yeast (yeast two-hybrid assay, Y2H) and *Agrobacterium*-mediated transient expression in plant tissue.
