## Supplemental Figures for "The core effector RipE1 of *Ralstonia solanacearum* interacts with and cleaves Exo70B1 and is recognized by the Ptr1 immune receptor"

### Supporting Information

**Figure S1:** Y2H negative controls, *Nicotiana sylvestris* HR assay

**Figure S2:** BiFC interaction between RipE1 mutants (C172A, ΔA) and Exo70B1.

#### Suppl. Figure S1

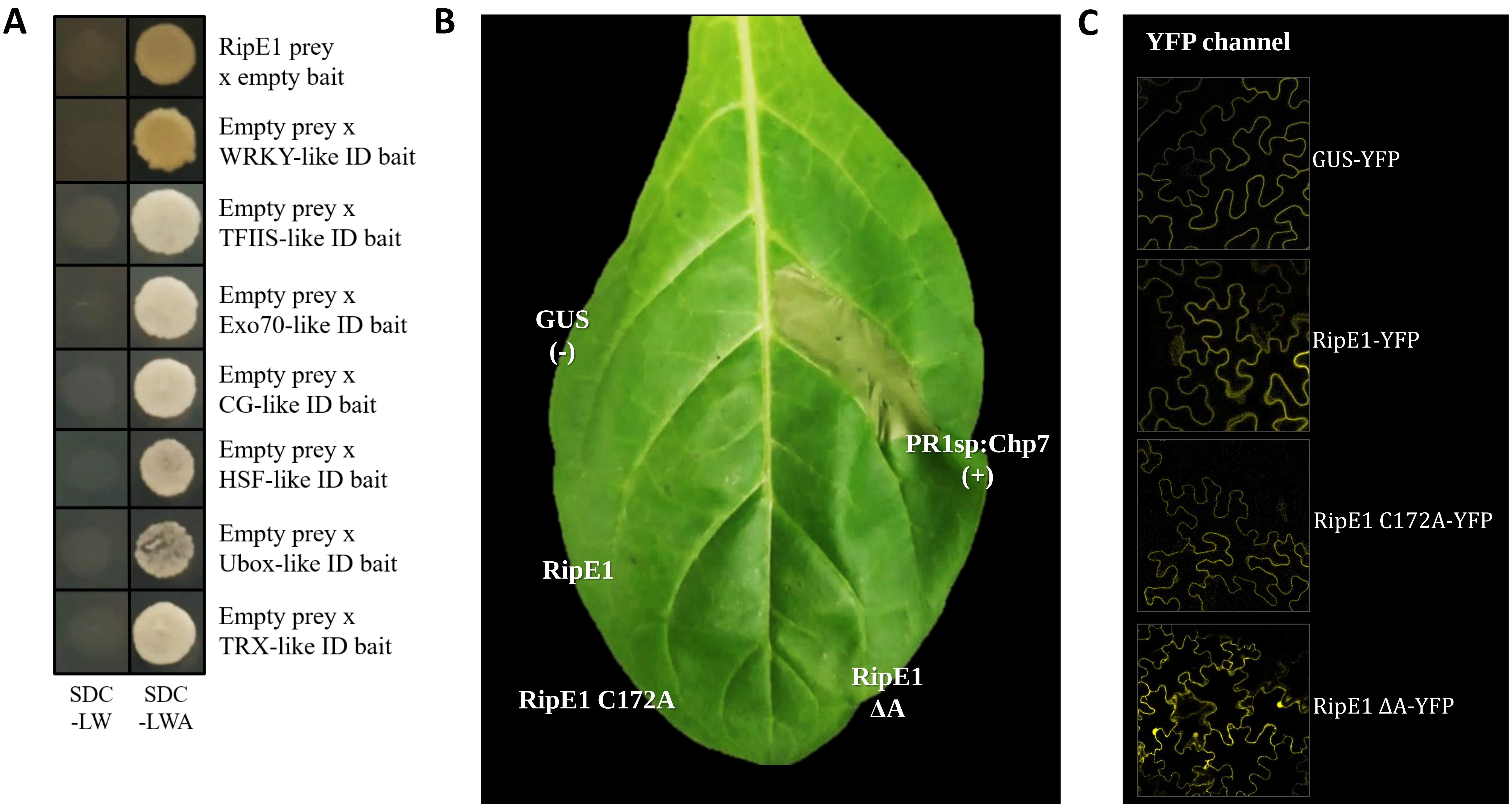


**Figure S1. Representative images supplementing main Figures 2 and 3. (A)** Neither RipE1 or any of the NLR-IDs were able to induce expression of the reporter gene in a yeast two-hybrid assay, in the absence of an interacting protein. Empty bait and prey vectors, respectively, were used to test possible auto-activities. The results support the interactions in yeast, that are presented in **Figure 2**. **(B)** Wild type RipE1 or mutants C172A and ΔA do not induce a hypersensitive response (HR) in *Nicotiana sylvestris* selected ecotype NS2706. Effector Chp7 –fused to PR1 secretion peptide for apoplastic secretion- was used as a positive HR-inducing control and GUS reporter gene was used as a negative control. This photograph was taken 4dpi. This experiment was repeated at least 6 times with the same exact results. **(C)** Leaf discs from **(B)** were observed via confocal microscopy to confirm expression of proteins of interest that did not induce HR. All indicated proteins were c-terminally fused to YFP fluorescent epitope. These images were captured 5dpi. **(B) and (C)** collectively support the further use of *Nicotiana sylvestris* NS2706 as a model for our *in-planta* experiments.

#### Suppl. Figure S2

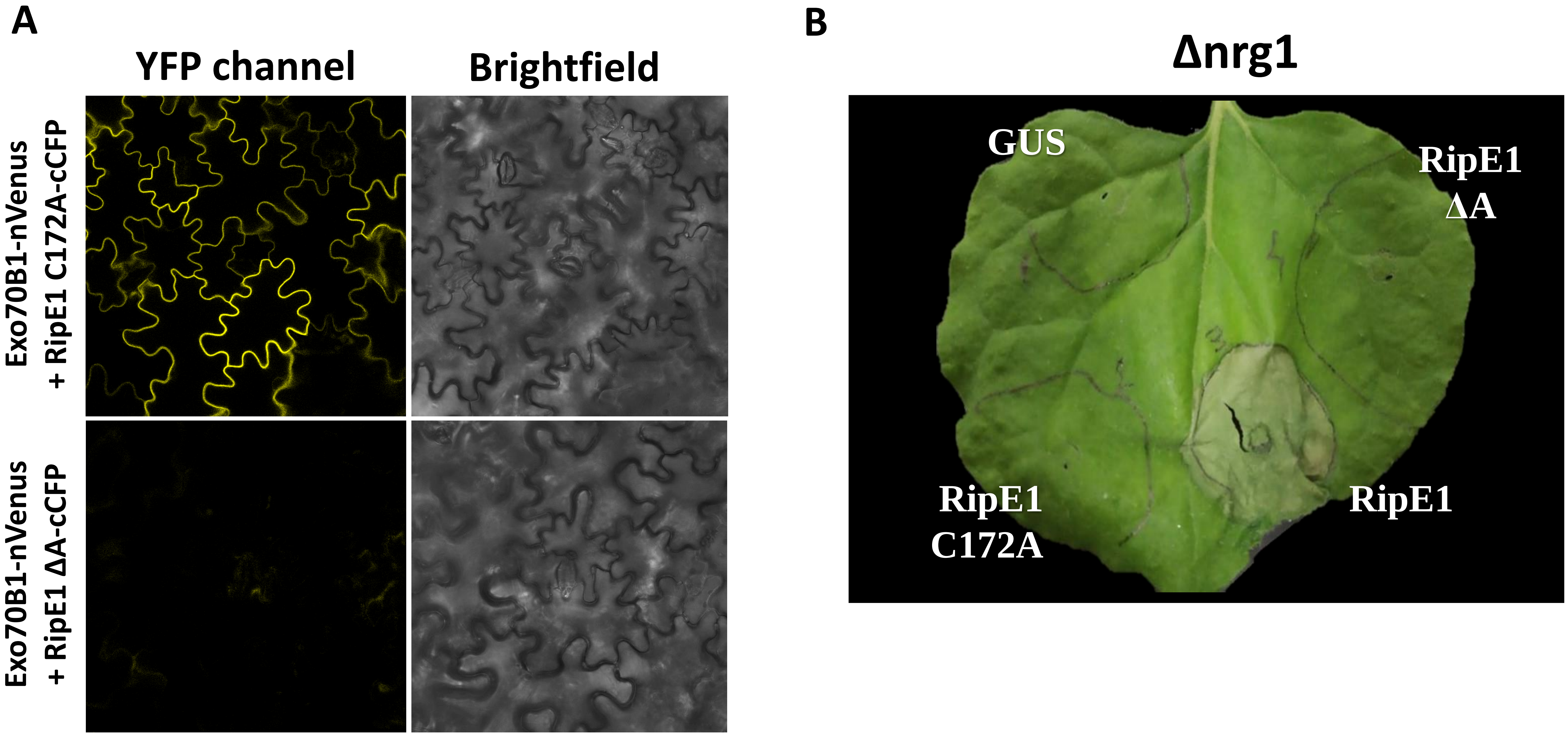


**Figure S2. (A)** Investigation of the RipE1-C172A/Exo70B1 and RipE1 ΔA/Exo70B1 association using a BiFC assay in *Nicotiana sylvestris* plants. RipE1 mutated proteins and Exo70B1 were fused at their C-termini with cCFP and nVenus epitope tags, respectively, and the YFP signal was detected via confocal microscopy 48hpi. YFP signal was detected when RipE1-C172A-cCFP was co-expressed with Exo70B1-nVenus, while it was not detected upon co-expression of RipE1 ΔA-cCFP/Exo70B1-nVenus. **(B)** RipE1-induced cell death in *N. benthamiana* is independent of NRG1. Wild type RipE1 and the mutants RipE1-C172A and RipE1 ΔA were transiently expressed in 4-week-old *N. benthamiana nrg1* plants via agroinfiltration. The experiment was performed 3 times with identical results. Photograph was taken 3dpi.

#### Suppl. Figure S3

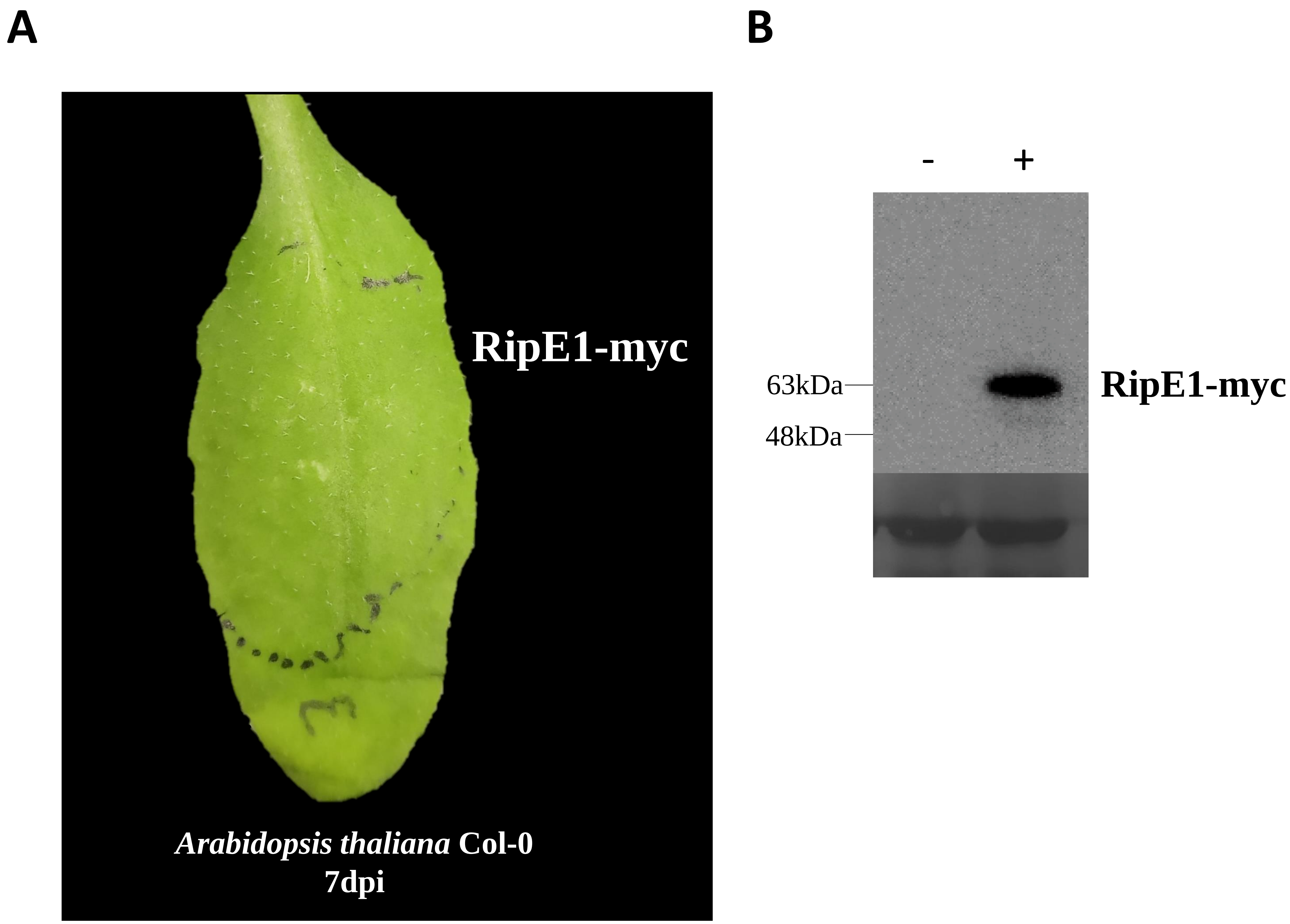


**Figure S3. RipE1 does not activate cell death after transient expression in Arabidopsis leaves. (A)** *Arabidopsis thaliana* Col-0 representative leaf, after transient expression of effector RipE1. The dotted circle marks the infiltrated area. RipE1-myc was transiently expressed via *Agrobacterium*-mediated infiltration at a total $OD_{600}$ of 0.5. Photographs were taken 7dpi. **(B)** RipE1-myc expression in (A) was confirmed via total protein extraction of both infiltrated (+) and non-infiltrated (-) areas and subsequent SDS-PAGE and western blot analysis.
